# An accurate molecular method to sex savannah elephants using PCR amplification of Amelogenin gene

**DOI:** 10.1101/856419

**Authors:** George G Lohay

**Affiliations:** Penn State University, 108 Life Sciences Building, Lee and Cavener Lab, University Park, PA, 16802, USA email: /

## Abstract

The use of molecular methods to identify the sex of elephants from non-invasive samples is essential for studies of population dynamics and population genetics. We designed a new technique for sex identification in savannah elephants using Amelogenin (AMEL) genes. The X-Y homologs of AMEL genes is known to be suitable for sex determination in pigs and some bovids. In this study on savannah elephants, the use of AMEL genes was more successful than previous methods that relied on genes found exclusively on Y-chromosomes, such as SRY, to distinguish males from females. We designed a common forward primer and two reverse primers for X- and Y-specific AMEL genes to obtain 262 bp and 196 bp PCR amplicons from X and Y genes, respectively. We tested the primers for the identification of the sex of 132 savannah elephants from fecal samples. The sex of 126 individuals (95.45%) matched observational data, while 6 (4.54%) did not match. This discrepancy observed was likely due to observational errors in the field, where high grass reduces the ability to accurately sex young individuals. Through our stool sample results, we have shown that the use of only three primers for AMELX/Y provides a highly accurate PCR-based method for sex identification in savannah elephants. The method is fast and shows more success than the SRY system by avoiding the inherent ambiguities of the previous PCR-based methods that made it difficult to distinguish between female samples and failed amplification reactions. Our sex identification method is non-invasive and can potentially be applied in population genetic studies and forensics tests on both savannah as well as forest elephants.

## Résumé

L’utilisation de méthodes moléculaires pour identifier le sexe des éléphants à partir d’échantillons non invasifs est essentielle pour les études de la dynamique et de la génétique des populations. Nous avons conçu une nouvelle technique d’identification du sexe chez les éléphants de savane à l’aide des gènes Amélogénine (AMEL). Les homologues X-Y des gènes AMEL sont connus pour être adaptés à la détermination du sexe chez les porcs et certains bovidés. Dans cette étude sur les éléphants de savane, l’utilisation des gènes AMEL a été plus efficace que les méthodes précédentes qui reposaient sur des gènes trouvés exclusivement sur les chromosomes Y, tels que SRY, pour distinguer les mâles des femelles. Nous avons conçu une amorce sens commune et deux amorces antisens pour les gènes AMEL spécifiques de X et Y pour obtenir des amplicons PCR de 262 pb et 196 pb à partir des gènes X et Y, respectivement. Nous avons testé les amorces pour l’identification du sexe de 132 éléphants de savane à partir d’échantillons de matière fécale. Le sexe de 126 individus (95,45%) correspondait aux données d’observation, tandis que 6 (4,54%) ne correspondaient pas. Cet écart observé était probablement dû à des erreurs d’observation sur le terrain, où les hautes herbes réduisent la capacité de déterminer le sexe des jeunes individus avec précision. Grâce aux résultats de nos échantillons de matière fécale, nous avons montré que l’utilisation de seulement trois amorces pour AMELX/Y fournit une méthode très précise basée sur la PCR pour l’identification du sexe chez les éléphants de savane. La méthode est rapide et montre plus de succès que le système SRY en évitant les ambiguïtés inhérentes aux précédentes méthodes basées sur la PCR qui rendaient difficile la distinction entre les échantillons femelles et les réactions d’amplification échouées. Notre méthode d’identification du sexe est non invasive et peut potentiellement être appliquée dans les études de génétique des populations et les tests médico-légaux sur les éléphants de savane et de forêt.

## Introduction

Molecular genetics is a powerful tool with many potential applications for conservation biology and wildlife management (Ortega et al. 2004). One of these applications includes the development of simple and accurate methods to determine the sex of wildlife from non-invasive samples (Strah and Kunej 2019). While poaching and illegal ivory trade are the most immediate threats to African savannah elephants, range and habitat fragmentation remain a significant longterm threat to the species’ survival (Douglas-Hamilton 1987; CITES 2014; Wittemyer et al. 2014). Poachers tend to kill elephant bulls with large tusks or adult females within families (Poole and Thomsen 1989). Poached elephant populations have been observed to have a skewed sex ratio in favor of females (Poole and Thomsen 1989; Aleper and Moe 2006; Jones et al. 2018). Traditional methods such as rapid demographic assessment (Poole 1989; Jones et al. 2018) which entirely rely on field observation of elephants to determine sex may produce errors due to the difficulty in distinguishing males from females in tall grass, especially among juveniles. An accurate method to sex elephants non-invasively is therefore crucial for many ecological studies, as well as for the identification of the forensic samples of elephant origin. For example, to assess age structure of elephants, it is important to determine whether the sex ratio for each class deviates from parity. Several methods have been developed that use PCR amplification of sexdetermination regions on chromosomes (Fernando and Melnick 2001; Gupta et al. 2006; Vidya et al. 2007; Ahlering et al. 2011). One method of sex identification for elephants that has been widely used on non-invasive sample types is based on three pairs of PCR primers for amplification of the genes AMELY2, SRY1, and PLP1 (Ahlering et al. 2011). The method requires conducting multiplex PCR, since females exhibit a single band (PLP1) and males exhibit all three bands, owing to the presence of Y-chromosome genes in males only. Although this method is reliable and more efficient than the previous methods (Ahlering et al. 2011), on some occasions it is hard to distinguish between a failed PCR and females, especially when male-specific primers did not work. Tests based on SRY amplification can be problematic as non-amplification of the Y-marker represents either female or a failed PCR (Peppin et al. 2010). An alternative method is based on the differences in sequence length between X- and Y-specific Amelogenin (AMEL) genes in a variety of species (Weikard et al. 2006). The X-Y homologs of AMEL genes (AMELX and AMELY) have been shown to be suitable for sex determination at the molecular level and have been used to identify sex in bovids (Weikard et al. 2006) and in pigs (Sembon et al. 2008). A single primer pair can be designed to amplify both chromosome-specific forms of the genes. The aim of this study was to design primers from AMEL genes specific for savannah elephants that can be used to accurately determine the sex of savannah elephants from non-invasive samples and provide researchers with additional resources. We tested our method by comparing the results using the technique developed by Ahlering et al. (2011).

## Materials and methods

We collected 132 fecal samples of savannah elephants from the Serengeti ecosystems in northern Tanzania. The fresh fecal samples were collected from external parts of dungs and stored in Queen’s College buffer solution buffer (20% DMSO, 0.25 M EDTA, 100 mM Tris, pH 7.5, saturated with NaCl). Four tissue samples (2 males and 2 females) from savannah elephants that died naturally were also included as positive controls for known sex. While in the field, an individual’s sex was determined through observations of genitalia, their overall body size, and the shape of the head and, in adults, the thickness of the tusks. Photographs were taken for each individual to confirm its sex. Through these closeup field observations, the sexes of all the individuals corresponding to each of the 132 samples were identified while collecting their fecal samples and these were used to test the new method. DNA was isolated using DNA extraction kits (QIAamp DNA Stool Mini Kit), following the manufacturer’s protocols with minor modifications (Eggert et al. 2005). Three PCR primers were designed using SnapGene^®^ software (GSL Biotech LLC), based upon the AMELX (Reference Sequence: NW_003573459) and AMELY (Genbank accession number AY823321) in *Loxodonta africana.* Three primers were designed: the common forward primer AMELXY 5’-TTCTGGAATCTGGTTTGAGGCT-3’, and the X-specific AMELX-R 5’-ATCTTTACAACAAAACAATTGTTAACCATGCTC-3’ and Y-specific AMELY-R 5’-TCAGATTCAGAGTTTCCTTCATGCAGTAG-3’ reverse primers.

Three primers would amplify both X and Y if present (i.e. males) yielding two DNA bands of different size, whereas females would generate only one band. The PCR reaction was performed with the initial polymerase activation step at 95°C for 3 min, denaturation at 95°C for 30 sec, annealing temperature at 56°C for 45 sec, extension at 72°C for 30 sec for 35 cycles. Each PCR mixture contained 3 μl of 5X Green GoTaq reaction buffer (promega), a final concentration of 0.67 μM of forward primers and 0.33 μM for each of the reverse primers, 0.13 μM of dNTP (Quanta bio), bovine serum albumin (BSA) 0.1 μg/μl, and 3 μl of DNA template of unknown concentration in a 15 μl volume. For each PCR reaction with test samples, we also ran a negative control (no DNA) and a positive control using DNA from known tissue samples. Finally, 6 μl of the PCR product was used for electrophoresis in Tris-Acetate EDTA running buffer at 120V for 45 min on a 2% agarose gel stained with GelRed (Biotium). Experiments were repeated twice to verify the results. Samples that gave ambiguous results were tested three times under the standard conditions described above. To confirm the origins of the PCR products, we sequenced amplicons from at least three samples for each primer pair and performed a BLASTn alignment search on the NCBI database to confirm the sequences of our primers. To validate our method, we compared our method with the method of Ahlering et al. (2011) by running PCR for 54 samples in parallel following the above procedure. We made this comparison to determine if our method could provide the same results as the previously published method.

## Results and discussion

A total of 132 dung samples were analyzed and successfully sexed using our AMELX/Y sexing method and, of these, 95.45% (126/132) samples were assigned to the gender corresponding to field observations (Table 1). In six cases (4.45%) the gender assigned using this PCR method did not match the gender identified in the field. Four of these were young elephants less than ten years of age. In the field, it is more difficult to accurately identify the sex of young elephants compared to older elephants due to limited visibility in long grass or savanna woodlands. A similar challenge was previously reported by Jones et al. (2017). Therefore, all mismatches observed were likely due to observational errors in the field and not the failure of PCR-based sexing method.

**Table 1.** The number of females and males identified using AMELX/Y primers, compared the reference sex based on field observations.

| Gender | Reference sex | AMELY/X sexing method | Mismatch |
| --- | --- | --- | --- |
| Females | 67 | 64 | 3 |
| Males | 65 | 62 | 3 |
| Total | 132 | 126 | 6 |

As expected, known male samples produced two DNA amplification bands whereas only one DNA band was amplified from female samples (fig. 1A). A similar method was used to identify the sex of bovids (Weikard, Pitra, and Kuhn 2006) and pigs (Sembon et al. 2008) using tissue samples. Because we used DNA derived from fecal samples, which have low amount of host DNA, we chose PCR priming sites that produced short fragment sizes of 262 bp and 196 bp for AMELX and AMELY respectively (fig. 1A).

**Figure 1:**
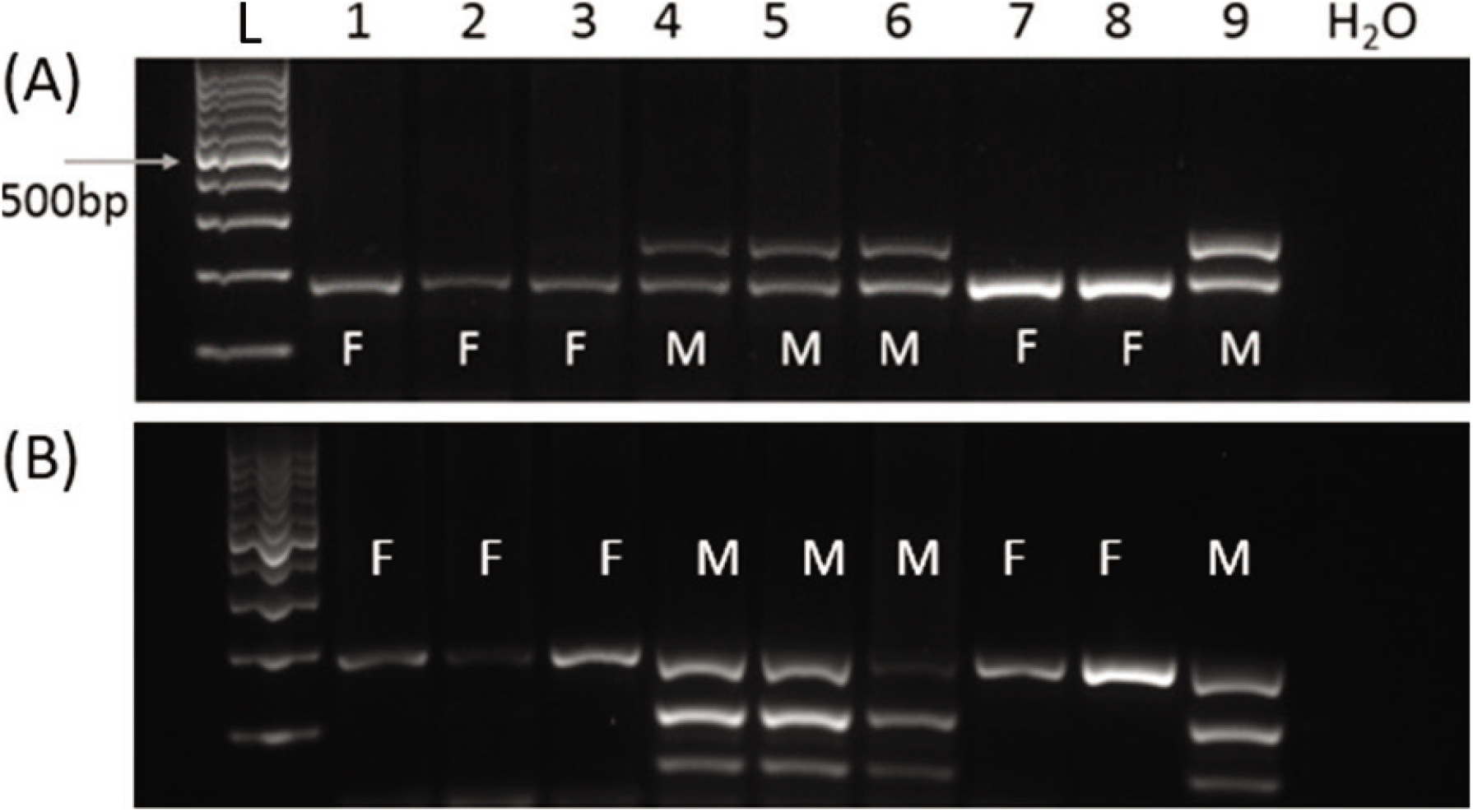
Agarose gel electrophoresis (2% agarose) of PCR product to determine sex of African savannah elephants. (A) AMELX and AMELY primers used in this study. Males (M) show two PCR bands (AMELX 196 bp and AMELY 262 bp) whereas females (F) show only one PCR band (AMELX 196 bp). Lane L is a100 bp DNA ladder (G-Biosciences) and water was used as a negative control. An arrow indicates the 500 bp mark on the ladder. (B) A gel image showing DNA bands using PLP1, AMELY2 and SRY1 (Ahlering et al. 2011), where males are identified by the presence of all three bands. Both methods (A) and (B) showed the same results using same samples under the same conditions.

PCR amplification was performed for 54 samples to compare our method and Ahlering et al. 2011 method. Our method had a lower PCR failure (2/54) rate than Ahlering et al. 2011 method (8/54). Except for two samples that could not amplify both Y-specific primers for males using Ahlering method (samples 11 and 37 in Table 2), the sex of all identified samples matched. We confirmed that our method works well with accuracy.

**Table 2.** Validation of AMELX/Y method using SRY system (Ahlering et al. 2011). M = male; F = females; M? = probably male; X = failed PCR. Differences in results between the two methods are indicated by italics.

| Sample ID | Ahlering et al. (2011) | This study | Sample ID | Ahlering et al. (2011) | This study |
| --- | --- | --- | --- | --- | --- |
| 1 | F | F | 28 | M | M |
| 2 | M | M | 29 | M | M |
| 3 | *X* | *M* | 30 | F | F |
| 4 | M | M | 31 | *M* | *X* |
| 5 | F | F | 32 | M | M |
| 6 | F | F | 33 | *X* | *M* |
| 7 | F | F | 34 | *M* | *X* |
| 8 | *X* | *F* | 35 | M | M |
| 9 | *X* | *F* | 36 | M | M |
| 10 | F | M | 37 | *M?* | *M* |
| 11 | *M?* | *M* | 38 | M | M |
| 12 | M | M | 39 | M | M |
| 13 | *X* | *M* | 43 | M | M |
| 14 | M | M | 41 | M | M |
| 15 | M | M | 42 | *X* | *M* |
| 16 | M | M | 43 | M | M |
| 17 | F | F | 44 | *X* | *M* |
| 18 | M | M | 45 | M | M |
| 19 | M | M | 46 | F | F |
| 20 | M | M | 47 | M | M |
| 21 | M | M | 48 | M | M |
| 22 | F | F | 49 | F | F |
| 23 | F | F | 50 | F | F |
| 24 | F | F | 51 | F | F |
| 25 | M | M | 52 | *X* | *M* |
| 26 | M | M | 53 | F | F |
| 27 | F | F | 54 | M | M |

Our method can be used as an alternative to the existing method (Ahlering et al. 2011) or the two methods can be run as parallel experiments to determine sex of savannah elephants (fig. 1). The advantage of AMELX/Y method is that there is no possibility of misidentifying a male as a female due to PCR failure. With our method PCR failure can be assessed by presence/absence of DNA bands. Whenever possible both methods should be applied to confirm the sex of all elephants from samples and clarify ambiguous results.

Using the previous multiplexing method of Ahlering et al. (2011), we unexpectedly found a higher number of failed PCR than our method (Table 2). A similar challenge was experienced in other studies which failed to identify the sex of between 16% and 23.8% of samples, due to repeated failure to discriminate DNA bands as either males or females (Moßbrucker et al. 2015; Lobora et al. 2017; Nofinska et al. 2019). Our method provides an additional resource for researchers interested on savannah elephant demography that reduces the risk of failure to accurately determine sex of individuals.

Conducting fieldwork is expensive and labor intensive, it is crucial to employ efficient methods to get as much data as possible. Missing data on the sex of individual elephants makes it more difficult to understand important aspects of population structure, such as effective population size. Our technique for identifying the sex of savannah elephants can be used in a wide variety of studies aimed at the conservation of savannah elephants, including those that address human–elephant conflicts, population dynamics, and anti-poaching efforts. Future improvements possible for this method, such as an automated assay, could further facilitate genetic analysis of free-ranging elephant populations and could make significant contributions to the management and conservation of other mammalian threatened species.

One of the limitations of our method is that the Y chromosome product is larger than the X product. The recommended length for use with degraded samples should be <170 bp (Strah & Kunej 2019, Durnin et al. 2007) or <250 bp according to (Villesen & Fredsted 2006). This could be problematic because in degraded samples the test described here may lead to males being incorrectly identified as females, when the Y band fails to amplify because it is has larger product length. We therefore advise that this test is used in combination with others like Ahlering et al. (2011) which has additional Y markers. Also, the common forward primer AMELXY can be redesigned to reduce the fragment size for Y.

In this article, we described a simple technique to sex savannah elephants using a non-invasive method. To our knowledge, AMELX/Y method has not been used before for sexing elephants based on DNA collected from blood/tissue or fecal samples in the field. This method could provide additional resources to researchers to accurately determine the sex of savannah elephants and other mammalian species as described in Weikard et al. (2006). We suggest further studies to be done on forest elephants using the PCR amplification of Amelogenin gene. This identification has potential to assist in forest environment studies for *L. a. cyclotis,* where it would be even more useful than for savannah elephants, which are more visible in their habitat.

## Acknowledgements

This work was funded by Cleveland Metroparks Zoo, Wildlife Conservation Society and the Huck Institutes of the Life Sciences. I thank Tanzania Wildlife Research Institute, Tanzania National Park Authority, and Tanzania Wildlife Authority for providing permissions to conduct research in protected areas. I thank my advisor Dr. Douglas Cavener for his assistance in designing the method and Dr. Anna Estes for her support during the fieldwork. I thank Thomas Schneyer for his assistance in the laboratory analysis, Chelsea Hudson, Jingjie Hu and Rebecca Bourne for their technical support and Dr. Barbara McGrath for proofreading of the manuscript.

## Notes

### Competing Interest Statement

The authors have declared no competing interest.

